# Inhibition of the replication of SARS-CoV-2 in human cells by the FDA-approved drug chlorpromazine

**DOI:** 10.1101/2020.05.05.079608

**Authors:** Marion Plaze, David Attali, Matthieu Prot, Anne-Cécile Petit, Michael Blatzer, Fabien Vinckier, Laurine Levillayer, Florent Perin-Dureau, Arnaud Cachia, Gérard Friedlander, Fabrice Chrétien, Etienne Simon-Loriere, Raphaël Gaillard

## Abstract

Urgent action is needed to fight the ongoing COVID-19 pandemic by reducing the number of infected people along with the infection contagiousness and severity. Chlorpromazine (CPZ), the prototype of typical antipsychotics from the phenothiazine group, is known to inhibit clathrin-mediated endocytosis and acts as an antiviral, in particular against SARS-CoV-1 and MERS-CoV. In this study, we describe the *in vitro* testing of CPZ against a SARS-CoV-2 isolate in monkey and human cells. We evidenced an antiviral activity against SARS-CoV-2 with an IC50 of ∼10μM. Because of its high biodistribution in lung, saliva and brain, such IC50 measured *in vitro* may translate to CPZ dosage used in clinical routine. This extrapolation is in line with our observations of a higher prevalence of symptomatic and severe forms of COVID-19 infections among health care professionals compared to patients in psychiatric wards. These preclinical findings support the repurposing of CPZ, a largely used drug with mild side effects, in COVID-19 treatment.

## To the Editor

From the beginning of the COVID-19 outbreak several weeks ago, we observed in Sainte Anne hospital (GHU PARIS Psychiatrie & Neurosciences, Paris, France) a higher prevalence of symptomatic and severe forms of COVID-19 infections among health care professionals (∼14%) compared to patients in psychiatric wards (∼4%) (1). This unexpected finding, that patients with higher comorbidities and risk factors (overweight, cardiovascular disorders…) seem to be protected against symptomatic and severe forms of COVID-19, drew our interest to decipher putative factors that could mediate this anti-SARS-CoV-2 protection. Because patients in psychiatric wards benefit from psychotropic medications, we screened the literature for antiviral effects associated with those drugs. This literature analysis identified chlorpromazine (CPZ), the prototype of phenothiazine-type antipsychotics, as the lead candidate (1). Indeed, CPZ has been widely used in clinical routine in the treatment of acute and chronic psychoses for decades. This first antipsychotic medication was discovered in 1952 by Jean Delay and Pierre Deniker at Sainte Anne hospital (2). In addition to its antipsychotic activity, a growing body of *in vitro* studies has demonstrated antiviral properties, for example against influenza (3), hepatitis viruses (4), alphaviruses (5), JC virus (6), Japanese encephalitis virus (7), bronchitis-virus (8), MHV-2 (9), Zika virus (10) or dengue virus (11). CPZ has also been shown to have antiviral activity against coronaviruses in multiple studies (12–14). It was identified active against both MERS-CoV and SARS-CoV-1 in a screen of 348 FDA-approved drugs, together with three other compounds (chloroquine, loperamide, lopinavir), using different cell lines (12). Similar results were obtained in a different library screen (13), as well as in another study using primary human monocytes (14). The genetic similarity between SARS-CoV-1 and SARS-CoV-2 suggests that this effect could also apply to this novel coronavirus. The antiviral activity of CPZ is mainly associated to inhibition of clathrin-mediated endocytosis (15–18), via translocation of clathrin and AP2 from the cell surface to intracellular endosomes (16). This clathrin-mediated endocytosis is essential for coronavirus cell entry (19). A very recent review article underlines the therapeutic potential of targeting clathrin-mediated endocytosis to tackle SARS-CoV-2 (20).

In this context, the aim of the current study was to investigate CPZ antiviral activity against SARS-CoV-2 in an *in vitro* assay. We therefore infected monkey VeroE6 cells with SARS-CoV-2 at a MOI of 0.1 for 2 h, in presence of different concentrations of CPZ. Supernatants were harvested at day 2 and analyzed by RT-qPCR for the presence of SARS-CoV-2 RNA (Figure 1.A). In parallel, cell viability was assessed on non-infected cells. While CPZ was associated with a cytotoxic effect in this model at the highest doses assessed, we measured an antiviral activity against SARS-CoV-2, with an IC50 of 9 μM. We also measured the viral RNA production in human A549-ACE2 cells (MOI of 1), where the cytotoxicity of CPZ was less pronounced (Figure 1.B), estimating an IC50 of 10.4 µM. These findings warrant further clinical investigations. Indeed, the biodistribution of CPZ (Figure 2) is highly compatible with the tropism of SARS-CoV-2: preclinical and clinical studies have reported high CPZ concentration in the lungs (20-200 times higher than in plasma (21–24)) and in saliva (30-100 times higher than in plasma (22,25)). Finally, CPZ readily crosses the blood-brain barrier (21–23,26) and could therefore prevent the neurological forms of COVID-19 (27).

**Figure 1.**
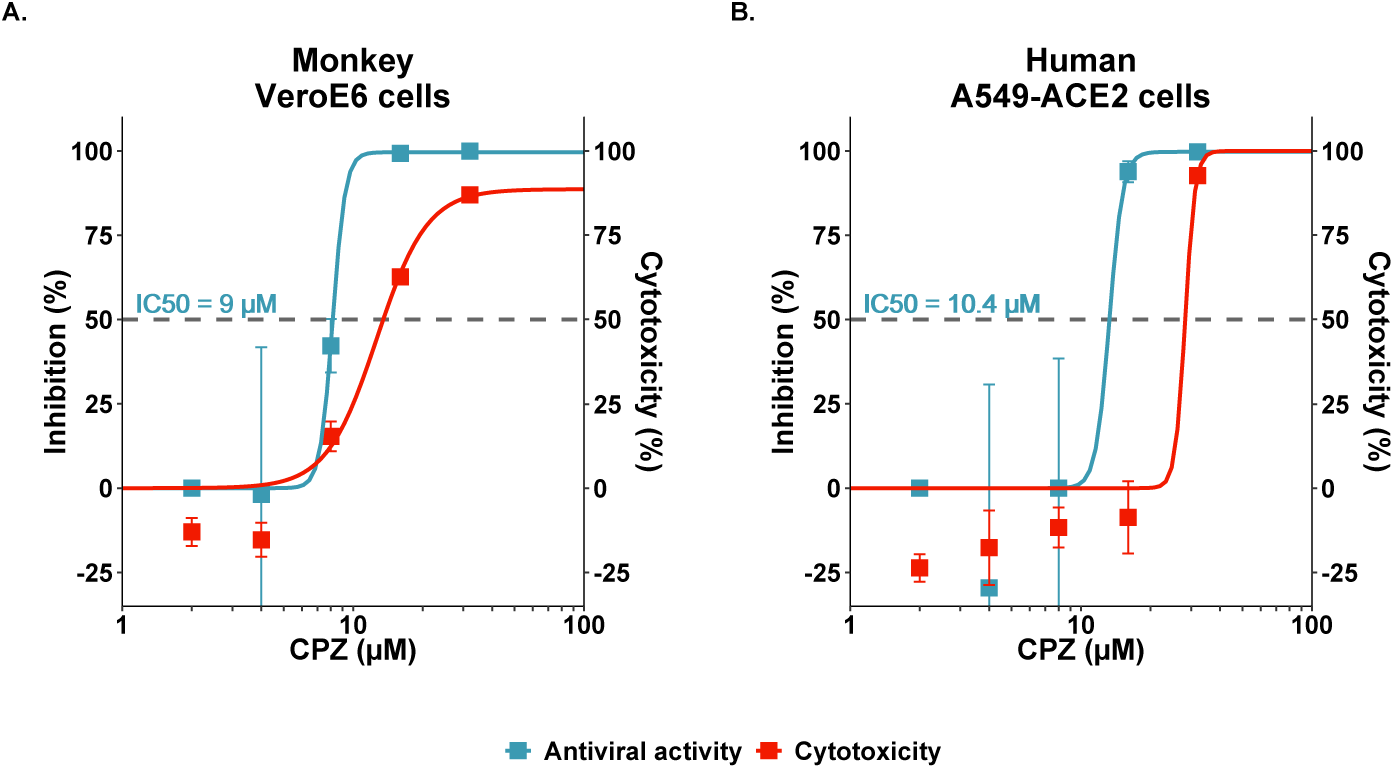
Antiviral activity of CPZ against SARS-CoV-2 in vitro in monkey VeroE6 cells (A) and human A549-ACE2 cells (B). Viral load in supernatants were measured at 48h (left Y axis), and viability under increasing concentrations of the antiviral compound are shown. Error bars denote s.e.m.

**Figure 2.**
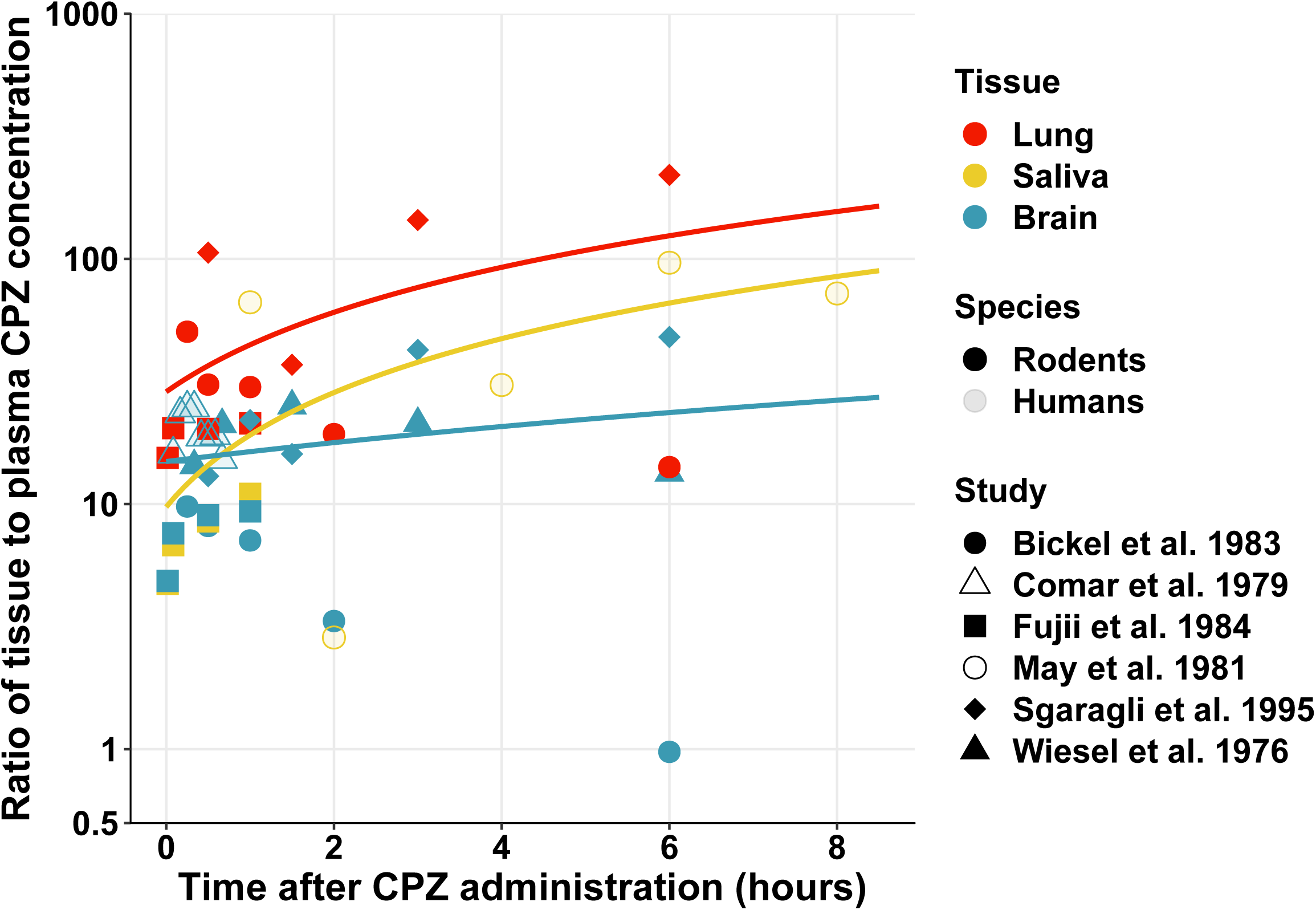
Review of temporal CPZ biodistribution in lung, saliva and brain. Ratio of tissue to plasma CPZ concentrations (log scale) after administration of a single dose of CPZ are represented for lung (red), saliva or salivary glands (yellow) and brain (blue) in rodents (filled) and humans (no-filled). Derived from previous preclinical and clinical studies (21–23,25,26,33).

With 2 900 000 infections and more than 200 000 deaths worldwide in just a few months (28), tools are urgently needed to help against the SARS-CoV-2 pandemic, to diminish disease severity along with contagiousness and to reduce the socio-economic consequences of the pandemic. Repurposing CPZ, a molecule already used in clinical practice, could offer both ready-to-use treatment with well-known and very mild side effects but also prophylactic strategy for the time after the lock down. At this time, CPZ is prescribed for around 70 years and FDA-approved in psychiatry and anesthesiology, with an excellent tolerance profile. CPZ is also used in clinical routine in nausea and vomiting of pregnancy (29), in advanced cancer (30), and to treat headaches in various neurological conditions (31). Even though the extrapolation from *in vitro* to clinically relevant dosage is not straightforward, the IC50 of 10μM (*i.e.* 3189 ng/ml; CPZ molar mass = 318.86 g/mol) measured *in vitro* may be compatible with CPZ dosage used in clinical routine. Indeed, residual plasma levels of CPZ in patients range from 30 to 300 ng/ml (32), which corresponds to 600 – 60 000 ng/ml in lungs (21–24) and 900 – 30 000 ng/ml in saliva (22,25). This extrapolation is supported by our observation of lower prevalence of symptomatic and severe forms of COVID-19 infections in psychiatric patients.

In conclusion, this first *in vitro* study of CPZ antiviral activity against SARS-CoV-2 in monkey and human cells supports the repurposing of CPZ, a well-known drug with antiviral properties and an excellent tolerance profile, to treat the actual COVID-19 pandemic for which effective treatments are still lacking. This proof of principle for the feasibility of CPZ as anti-SARS-CoV-2 therapeutic is a critical step for ongoing clinical trial (NCT04366739).

## Material & Methods

### Cell culture and virus isolates

Vero E6 cells (African green monkey kidney epithelial cells, ATCC, CRL-1586) were maintained in Dulbecco’s modified Eagle’s medium (DMEM) containing 10% fetal bovine serum (FBS) and 5 units/mL penicillin and 5 μg/mL streptomycin at 37°C with 5% CO2. A549-ACE2 cells (adenocarcinomic human alveolar basal epithelial cells, transduced to express the human Angiotensin-converting enzyme 2 (ACE2), kind gift of Pr. O. Schwartz, Institut Pasteur, Paris France) were maintained in DMEM containing 10% FBS, 5 units/mL penicillin and 5 μg/mL streptomycin and 40 µg/mL blasticidin at 37°C with 5% CO2.

SARS-CoV-2, isolate BetaCoV/France/IDF0372/2020 C2, was supplied by the National Reference Centre for Respiratory Viruses (NRC) hosted at Institut Pasteur (Paris, France) and headed by Pr. S. Van der Werf. The human sample from which this strain was isolated has been provided by Dr. X. Lescure and Pr. Y. Yazdanpanah from the Bichat Hospital, Paris, France. Viral stocks were prepared by propagation in VeroE6 cells in DMEM supplemented with 2% FBS. Viral titers were determined by plaque assay. All experiments involving live SARS-CoV-2 were performed in compliance with Institut Pasteur guidelines for Biosafety Level 3 work. All experiments were performed in at least three biologically independent replicates.

### Antiviral activity assay

Cells were seeded into 96-well plates 24 h prior to the experiment. Two hours prior to infection, cell culture supernatant was replaced with media containing 32µM, 16µM, 8µM, 4µM and 2µM of CPZ, or the equivalent volume of maximum H_2_O vehicle used as a control. For the infection, the drug-containing media was removed, and replaced with virus inoculum (MOI of 0.1 PFU/cell for VeroE6 and 1 for A549-ACE2) for 2 hours. The inoculum was then removed and replaced with 100µl fresh media (2% FBS) containing CPZ at the indicated concentrations or H_2_O and incubated for 48 hours.

At 48h, cell supernatant was collected and spun for 5 min at 3,000g to remove debris. Toxicity controls were setup in parallel on uninfected cells.

RNA was extracted from 50µl aliquots of supernatant using the Nucleospin 96 virus kit (Macherey-Nagel) following the manufacturer’s instructions. Detection of viral genomes was performed by RT-qPCR, using the IP4 primer set developed by the NRC at Institut Pasteur (described on the WHO website (https://www.who.int/docs/default-source/coronaviruse/real-time-rt-pcr-assays-for-the-detection-of-sars-cov-2-institut-pasteur-paris.pdf?sfvrsn=3662fcb6_2). RT-qPCR was performed using the Luna Universal One-Step RT-qPCR Kit (NEB) in an Applied Biosystems QuantStudio 3 thermocycler, using the following cycling conditions: 55°C for 10 min, 95°C for 1 min, and 40 cycles of 95°C for 10 sec, followed by 60°C for 1 min. The quantity of viral genomes is expressed as PFU equivalents, and was calculated by performing a standard curve with RNA derived from a viral stock with a known viral titer. IC50 values were fitted using four-parameter dose response curves in GraphPad prism v8.4.2.

Cell viability in drug-treated cells was measured using the AlamarBlue reagent (ThermoFisher). At 48 h post treatment, the drug-containing media was removed and replaced with AlamarBlue and incubated for 2h at 37°C. Fluorescence was measured in a Tecan Infinity 2000 plate reader. Percentage viability was calculated relative to untreated cells (100% viability).

## Acknowledgments

We are grateful to the Centre National de Reference des virus des infections respiratoires for sharing reagents and protocols. We thank Olivier Schwartz and his team for sharing the A549-ACE2 cell line. This study has received funding from Institut Pasteur (covid-therap) and the French Government’s Investissement d’Avenir program, Laboratoire d’Excellence “Integrative Biology of Emerging Infectious Diseases” (grant n°ANR-10-LABX-62-IBEID). ESL acknowledges funding from the INCEPTION program (Investissements d’Avenir grant ANR-16-CONV-0005).

